# Shear and hydrostatic stress regulate fetal heart valve remodeling through YAP-mediated mechanotransduction

**DOI:** 10.1101/2022.11.24.517814

**Authors:** Mingkun Wang, Belle Yanyu Lin, Shuofei Sun, Charles Dai, FeiFei Long, Jonathan T. Butcher

## Abstract

Clinically serious congenital heart valve defects arise from improper growth and remodeling of endocardial cushions into leaflets. Genetic mutations have been extensively studied but explain less than 20% of cases. Mechanical forces generated by beating hearts drive valve development, but how these forces collectively determine valve growth and remodeling remains incompletely understood. Here we decouple the influence of those forces on valve size and shape, and study the role of YAP pathway in determining the size and shape. The low oscillatory shear stress promotes YAP nuclear translocation in valvular endothelial cells (VEC), while the high unidirectional shear stress restricts YAP in cytoplasm. The hydrostatic compressive stress activated YAP in valvular interstitial cells (VIC), whereas the tensile stress deactivated YAP. YAP activation by small molecules promoted VIC proliferation and increased valve size. YAP inhibition suppressed the VIC proliferation and reduced valve size, but enhanced cell-cell adhesions between VEC thus maintaining an elongated shape. Finally, left atrial ligation was performed in chick embryonic hearts to manipulate the shear and hydrostatic stress in-vivo. The restricted flow in the left ventricle induced a globular and hypoplastic left atrioventricular (AV) valves with an inhibited YAP expression. By contrast, the right AV valves with sustained YAP expression grew and elongated normally. This study establishes a simple yet elegant mechanobiological system by which transduction of local stresses regulates valve growth and remodeling. This system guides leaflets to grow into proper sizes and shapes with the ventricular development, without the need of a genetically prescribed timing mechanism.

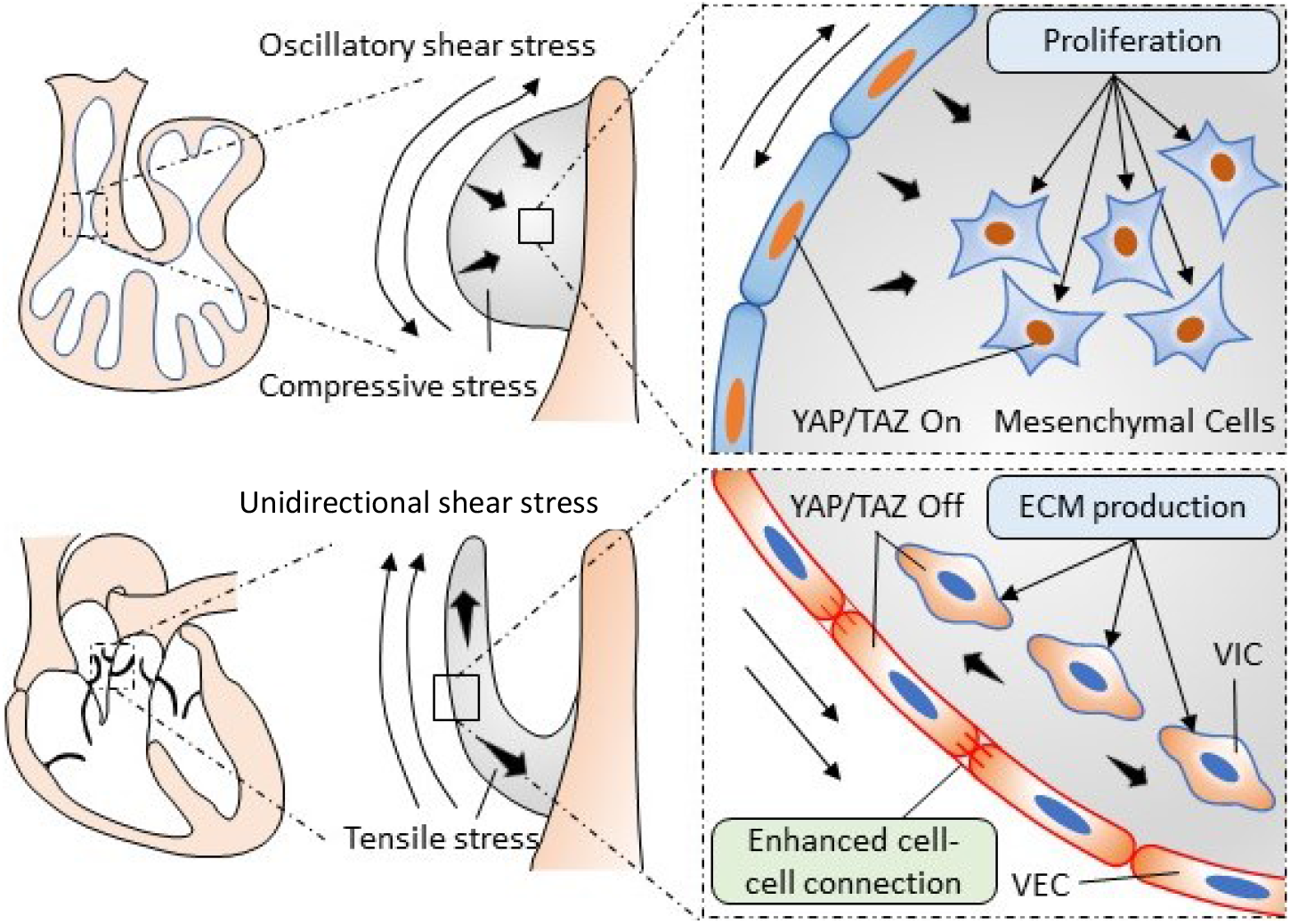

## Introduction

Congenital heart disease is the most common birth defects and affects about 0.5 – 2.0% of the general population.(1) Congenital heart valve defects accounts for over 25% of all congenital heart disease.(2) They can be immediately life threatening at birth or impair the long-term cardiac function in adulthood.(3) Heart valve development starts with endothelial-mesenchymal-transition (EMT), in which valvular endothelial cells (VEC) gain mesenchymal markers and invade into the subendothelial matrix to form endocardial cushions. Post-EMT, these cells proliferate and differentiate into extracellular matrix (ECM) producing valvular interstitial cells (VIC). With precise regulation of VEC and VIC, the cellularized endocardial cushions undergo ECM remodeling and elongate into thin mature leaflets or cusps. During this process, disturbed growth and remodeling will result in valve malformation and cause clinically relevant cardiac defects.(4-6) Genetic causes of this disturbance have been extensively studied,(7-9) but can explain less than 20% of clinical cases.(10-12) The importance of mechanical forces in regulating valve development has become well appreciated.(13-15) Oscillatory shear stress (OSS) promotes EMT, and its cellular and molecular mechanisms have been well understood.(16) By contrast, the role of mechanical forces in clinically important post-EMT growth and remodeling remain poorly understood.

The flowing blood generates shear and hydrostatic stress on valves. The shear stress is in direct contact with VECs, while the hydrostatic stress causes compression and tension in valves and can be transmitted to VICs. Multiple mechanosensitive signaling pathways have been identified in adult tissues,(17-19) but their involvement in valve morphogenesis is unclear. Although it has been shown in zebrafish that shear stress regulates early valvulogenesis (namely EndMT) via bioelectric signaling,(20) the near universal VEC in zebrafish heart valves limits study of these later growth and remodeling phases.(21) The contributions of VICs and hydrostatic stress, including potential collaborations with VEC, in valvulogenesis are not known. YAP signaling is another widely investigated mechanoactive pathway,(22, 23) and transcriptional cofactor of the Hippo pathway.(24) Yap signaling is known to regulate cardiac ventricular development by promoting cardiomyocyte proliferation.(25) However, its role in post-EndMT valvular morphogenesis is poorly understood. Moreover, whether YAP pathway responds differently to different forces in different cell types, and how these mechanoresponses collaboratively regulate multiple cell types for a specific tissue morphogenesis, are not known.

Here we explored the mechanism by which the shear and hydrostatic stress regulate valve growth and remodeling. We used in-vitro and ex-vivo models to decouple the effects of shear and hydrostatic stress on the size and shape of valves. We also studied the role of YAP mediated mechanotransduction in those effects by gain- and loss-of-function tests. To verify our findings in-vivo, we performed left atrial ligation (LAL) in chick embryonic hearts. The four-chambered chick hearts develop in a manner that mirrors human heart development. The LAL manipulates mechanical forces in-vivo and has been shown to replicate some important features of congenital heart defects.(5, 26, 27) By combining those models, we elaborate how shear and hydrostatic stress regulates VEC and VIC to determine proper size and shape for valves.

## Results

### YAP expression is spatiotemporally regulated

We collected embryonic hearts at different development stages from wild type mice and examined the expression pattern of YAP in heart valves. We found that YAP was expressed in both mesenchyme and endothelium of outflow tract (OFT) and atrioventricular (AV) cushions at E11.5 (Figure 1A). YAP expression in VIC increased significantly at E14.5 (Figure 1B) then dropped significantly at E17.5 (Figure 1C). Whereas YAP expression in VECs decreased at E14.5 and further decreased at E17.5. The expression followed the same pattern in both semilunar (SL) and AV valves (Figure 1D). At E17.5, YAP expression in VICs localized in or near tips of SL and AV valves (dashed circles), while in VECs, YAP was mainly expressed on fibrosa (solid arrows) and absent on the ventricularis of SL valves and atrialis of AV valves (dashed arrows).

**Figure 1.**
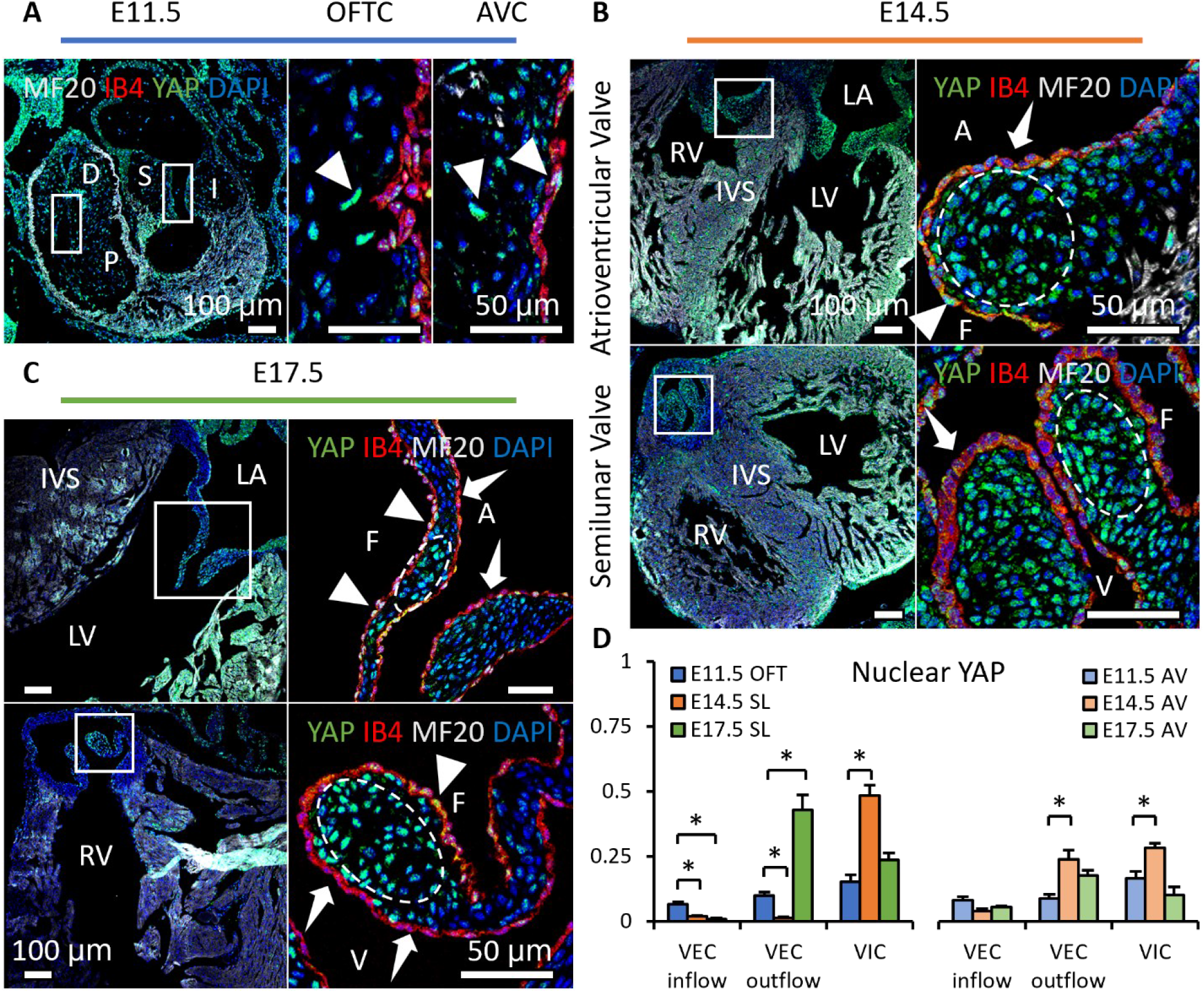
YAP expression is spatiotemporally regulated and correlated to mechanical environments during valve remodeling. **A-C**. Heart sections from E11.5, E14.5, E17.5 were performed immunofluorescence staining with YAP (Green), endothelial maker IB4 (red), myocardial marker MF20 (gray) and DAPI (blue). **A**. YAP expresses in both cushion mesenchyme and endothelium (solid arrows) at E11.5. **B**. YAP is intensively expressed in VICs (white circles) at E14.5 but seldom detected in VECs (dashed arrows). **C**. YAP expression remains in VICs on the tip regions and in VECs on the fibrosa side at E17.5 (solid arrows), while disappears in VECs on the ventricularis or ventricularis (dashed arrows) **D**. Ratios of VECs and VICs expressing nuclear YAP in AV and SL valves at different stages, data are presented by mean ± SEM, n = 6 sections from 3 embryos, *p < 0.05, two-way ANOVA tests. OFTC, outflow tract cushion; AVC, atrioventricular cushion; I, inferior cushion; S, superior cushion; D, distal cushion; P, proximal cushion; LA, left atrium; LV, left ventricle; RV, right ventricle; IVS, interventricular septum; V, ventricularis; A, atrialis; F, fibrosa.

This side specific expression of YAP in VECs correlated with the different flow patterns: unidirectional shear stress (USS) develops on the outflow side, while OSS develops on the inflow side (Figure 3A). The localization of YAP positive VICs valve tips also correlated with the local compressive stress (CS) developed in the tips (Figure 3B).

### YAP co-expressed with LEF1 during valve remodeling

LEF1 is a transcriptional co-factor of WNT/ β-catenin signaling pathway, which promotes valve growth.(28) However, recent studies showed a discrepancy between the LEF1 and β-catenin expression in mouse embryonic heart valves, indicating that the pro-growth role of LEF1 is independent of β-catenin.(29) Here we found an extensive co-expression of LEF1 and YAP during valve remodeling. At E14.5, more than 75% of YAP expressing VIC also expressed LEF1 in both AV and OFT SL valves (Figure 2B). At E17.5, both YAP and LEF1 expressed were localized in the tip of valves (Figure 2C), but LEF1 expression in AV valves weakened. By contrast, at E11.5, no overlap between LEF1 and YAP was detected (Figure 2A). LEF1 was exclusively expressed in the subendothelial region, but YAP expression appeared to be arbitrary. This supports that YAP has different functions at early EndMT than later remodeling stages. The co-expression of LEF1 and YAP in VICs at later stages shows a pro-growth role of YAP in VICs.

**Figure 2.**
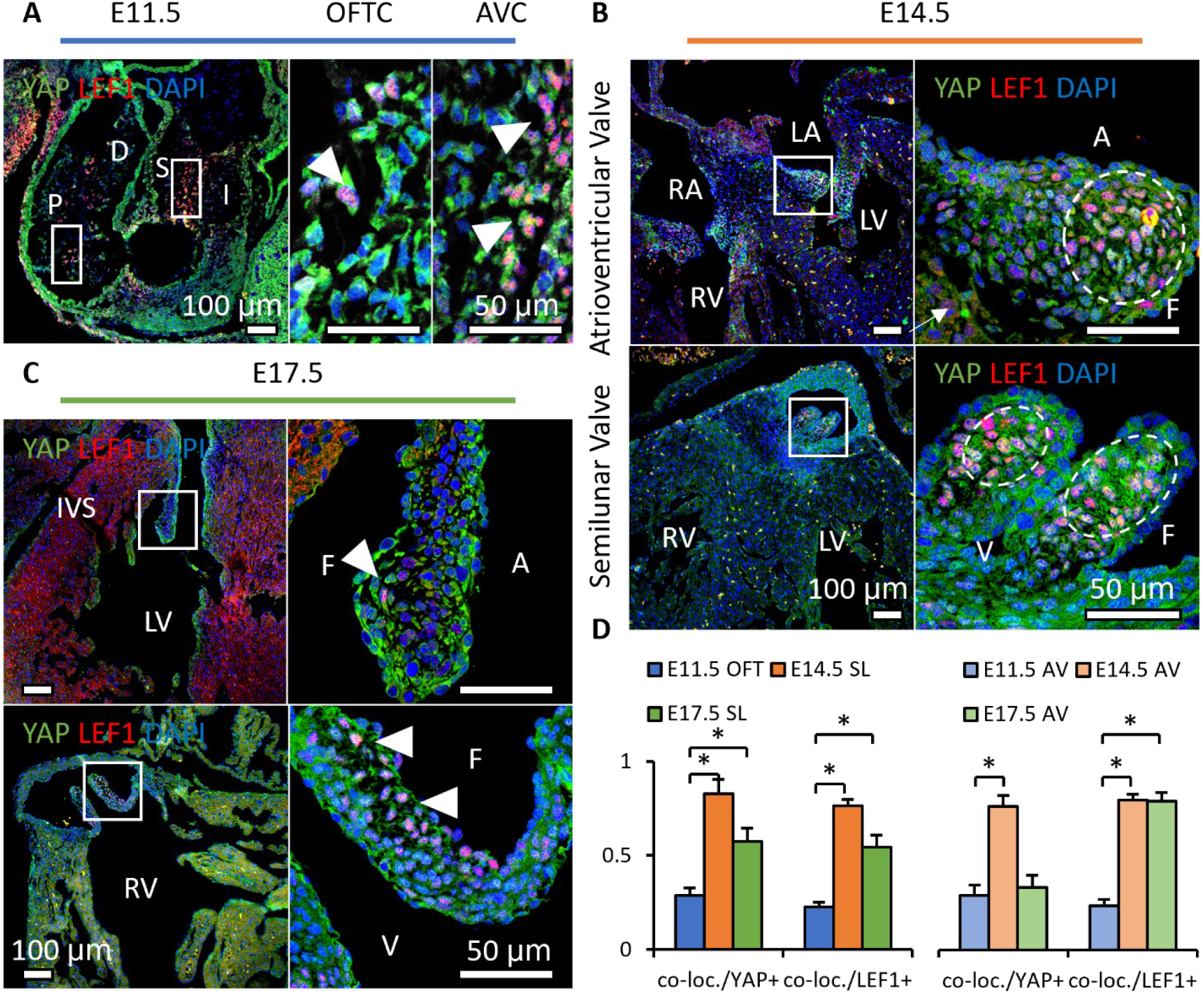
Nuclear YAP is associated with the growth phenotype in valves during remodeling. **A-B**. Co-expression of YAP and LEF1 at E11.5, E14.5, E17.5 by immunofluorescence staining with YAP (Green), LEF1 (red) and DAPI (blue), arrows show co-localization of YAP and LEF1. **A**. YAP and LEF1 expressions are not mutually exclusive at E11.5. **B**. YAP and LEF1 are co-expressed in the same or adjacent VICs at E14.5 (circles). **C**. YAP and LEF1 are co-localized in the VICs on the valve tip (arrows). **D**. Ratios of co-localized expression in all cell populations, YAP+ cells and LEF1+ cells of AV and SL valves at different stages. OFTC, outflow tract cushion; AVC, atrioventricular cushion; I, inferior cushion; S, superior cushion; D, distal cushion; P, proximal cushion; LA, left atrium; RA, right atrium; LV, left ventricle; RV, right ventricle; IVS, interventricular septum; V, ventricularis; A, atrialis; F, fibrosa. Scale Bar = 50μm. *p < 0.05

### Shear and hydrostatic stress regulate YAP activity

Since the spatiotemporal expression of YAP correlated with the pattern of shear stress and the direction of hydrostatic stress, we used in-vitro and ex-vivo tests to decouple the effects of shear and hydrostatic stress on the YAP signaling pathway in VECs and VICs.

To study the effect of shear stress on the YAP activity in VECs, we applied USS and OSS directly onto a monolayer of freshly isolated VECs. The VEC was obtained from AV cushions of chick embryonic hearts at Hamburger–Hamilton stage (HH) 25. The cushions were placed on collagen gels with endocardium adherent to the collagen and incubated to enable the VECs to migrate onto the gel. We then removed the cushions and immediately applied the shear flow to the monolayer for 24 hours. The low stress OSS (2 dyn/cm^2^) promoted YAP nuclear translocation in VEC (Figure 3C), while high stress USS (20 dyn/cm^2^) restrained YAP in cytoplasm.

**Figure 3.**
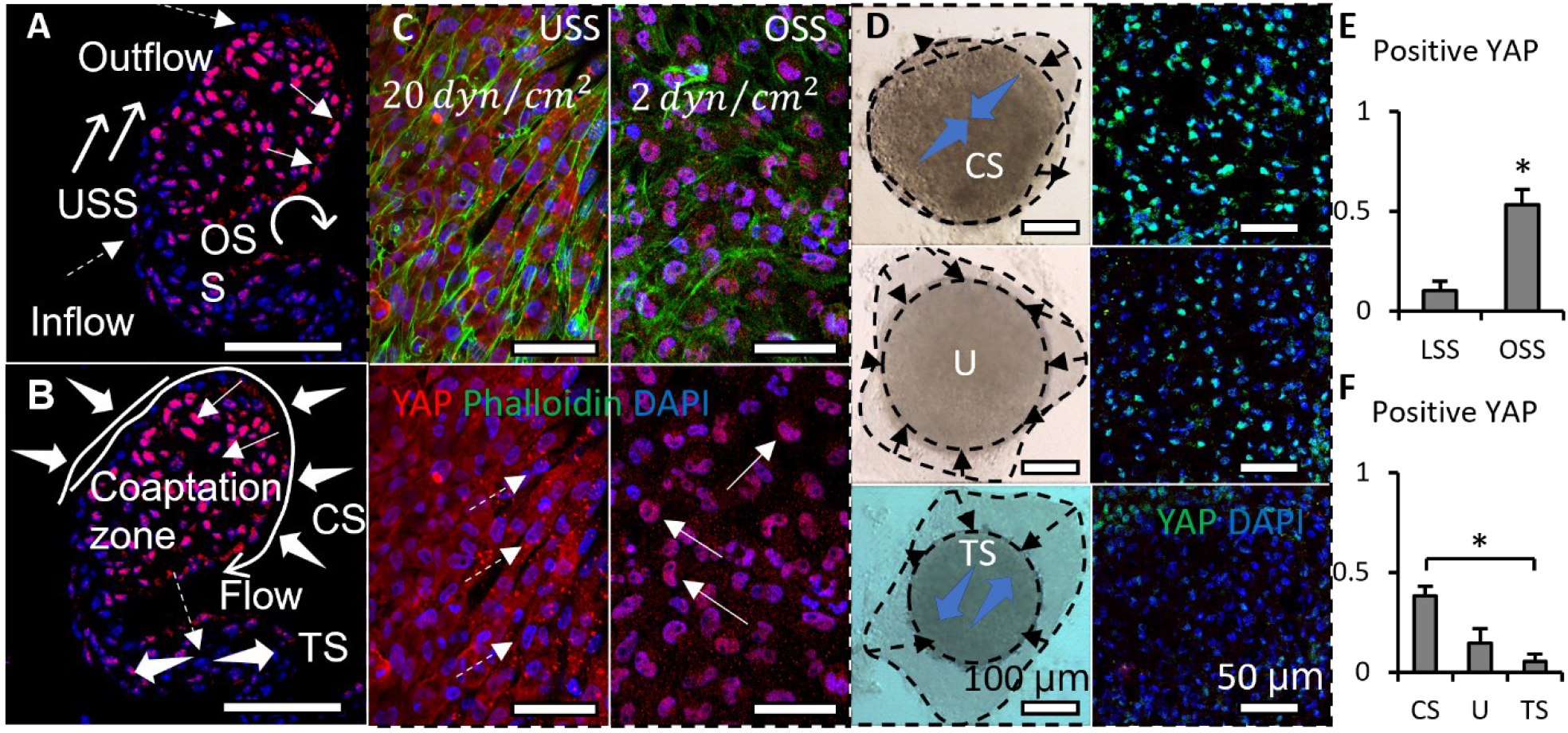
Shear stress and hydrostatic pressure regulate YAP activity. **A**. Illustration of shear flow distribution around a E17.5 pulmonary valve, and its correlation with the side-specific expression of YAP in VEC (positive: solid arrows; negative: dashed arrows) YAP. **B**. Illustration of compressive and tensile stress applied on a E17.5 pulmonary valve, and its correlation with the location-dependent expression of YAP in VIC (positive: solid arrows; negative: dashed arrows). **C**. YAP expression in a monolayer of VEC exposed to high stress (20 dyn/cm^2^) unidirectional flow, low stress (2 dyn/cm^2^) oscillatory flow. YAP, red; F-actin, green; DNA, blue. **D**. Bright field and immunofluorescence images (YAP, green; DAPI, blue) of cushion explants cultured under compress stress (CS), tensile stress (TS) and unloaded (U) conditions for 24 hours. The bright field images show the morphologies (dashed circles) and morphological changes (dashed arrows) of cushion explants, blue arrows represent the force. **E**. Influence of LSS and OSS on YAP activation in the monolayer VEC. **F**. Ratios of cells expressing YAP under different loading conditions. Data are presented by mean ± SEM, n = 15 explant valves from 8 embryos, *p < 0.05, two-way ANOVA tests or two-tailed student t-tests

To study the effect of hydrostatic stress on the YAP activation in VICs, we used media with different osmolarities to mimic the CS and tensile stress (TS).(30) CS was induced by hypertonic condition while TS was created by hypotonic condition, and unloaded (U) condition refers to the osmotically balanced media. We cultured HH25 AV cushion explants under those conditions for 24 hours and found that the trapezoidal cushions adopted a spherical shape (Figure 3D). TS loaded cushions significantly compacted, and the YAP expression in VICs of TS loaded cushions was significantly lower than that in CS loaded VICs (Figure 3F).

### Loss of YAP limited cell proliferation and led to a folded endothelium

To study the function of YAP in valve growth and remodeling, we added a pharmacological inhibitor of YAP, verteporfin (VP), into the CS (pro-growth) and U conditions. The VP inhibits the interaction between YAP and TEAD, which in turn, blocks transcriptional activation of targets downstream of YAP.(31) We isolated HH34 OFT SL valve explants and cultured them under CS, CS+VP and U+VP. We confirmed that the concentration of VP we used (5mM) does not influence cell viability (Figure S2). Successful YAP inhibition reduced the valve size (Figure 4A vs. Figure 4B), regardless of the media condition. In contrast to the spherical shape of valves cultured under CS, valves cultured in media with VP maintained their trapezoidal shape, which was characterized by circularity (Figure 3E). Further investigation demonstrated that loss of YAP significantly reduces the expression of the proliferation maker, pHH3 in the VIC (arrows, Figure 4C).

**Figure 4.**
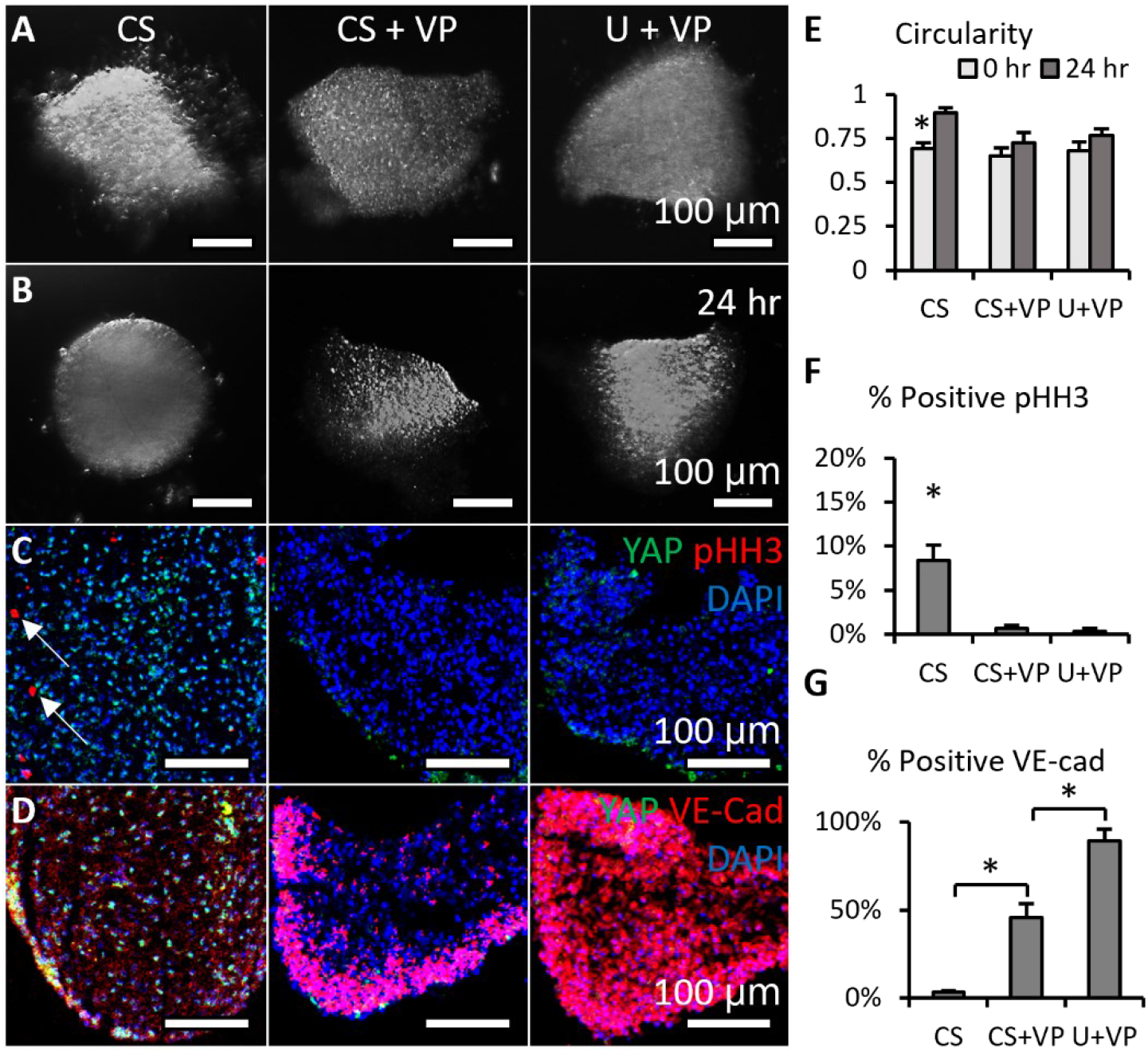
Loss of YAP limited cell proliferation and maintained a valvular shape. **A-B**. Bright field images of HH34 SL valve explants cultured under CS (compressive stress), CS+VP (compressive stress + YAP inhibitor), U+VP (unloaded + YAP inhibitor) conditions for 24 hours. **C**. After the 24-hr-culture, sections of explants were stained for YAP (green) and proliferation marker pHH3 (red, arrows), **D**. or YAP (green) and endothelial cell-cell junction VE-Cadherin (red). **E**. Comparison of explant morphology before and after 24 hour-culture. **F**. Comparison of percentages of cells expressing pHH3 under different culture conditions. **G**. Comparison of signal intensities of VE-Cad expression under different culture conditions, the intensities are normalized to maximum intensity. Data are presented by mean ± SEM, n = 15 explant valves from 8 embryos, *p < 0.05, two-way ANOVA tests or two-tailed student t-tests

In addition, the YAP inhibition significantly strengthened the expression of VE-cadherin between VECs (Figure 4D). The endothelium was normally a single layer of VECs, but the endothelium with strong VE-cad expression consisted of five layers of tightly packed VECs. The extra-thick layers could be a result of endothelium folding to minimize surface tensions.(32) The diameters of VP treated cushions D_VP_ ∼ 0.4 – 0.5 x D_0_ (diameters before VP treatment), which gives the surface area after the treatment A_VP_ ∼ 0.16 – 0.25 x A_0_ (before treatment), since the VEC number remained the same (survival rate > 90%) or increased (if the VEC proliferated), there must be more (1/ A_VP_ ∼ 4 – 5) layers of VECs. This estimation showed that loss of YAP led to endothelium folding.

### Activation of YAP promoted cell proliferation and led to a relaxed endothelium

For YAP gain-of-function study, a small molecule PY-60 was added into TS (pro-compaction) and U conditions. The PY-60 treatment dose-dependently promoted the association of YAP and TEAD. Unlike most YAP activators, PY-60 does not influence the kinases involved in mechanosensing.(33) PY-60 did not affect cell survival rate (Figure S2). YAP activation reversed the compaction of valves under TS and U conditions and drove a growth in valve size (Figure 5A vs. Figure 5B). All valves adopted the spherical shape no matter whether they grew or compacted. The pHH3 staining showed that the YAP activation increased VIC proliferation even under the TS condition (arrows, Figure 5C). In contrast to the multilayer structure in YAP depleted endothelium, the YAP expressing endothelium was single-layer and free of shrinkage (Figure 5D). The expression of VE-cadherin in the YAP activated endothelium was significantly weaker than that in YAP inhibited endothelium, showing that YAP activation led to a relaxed endothelium.

**Figure 5.**
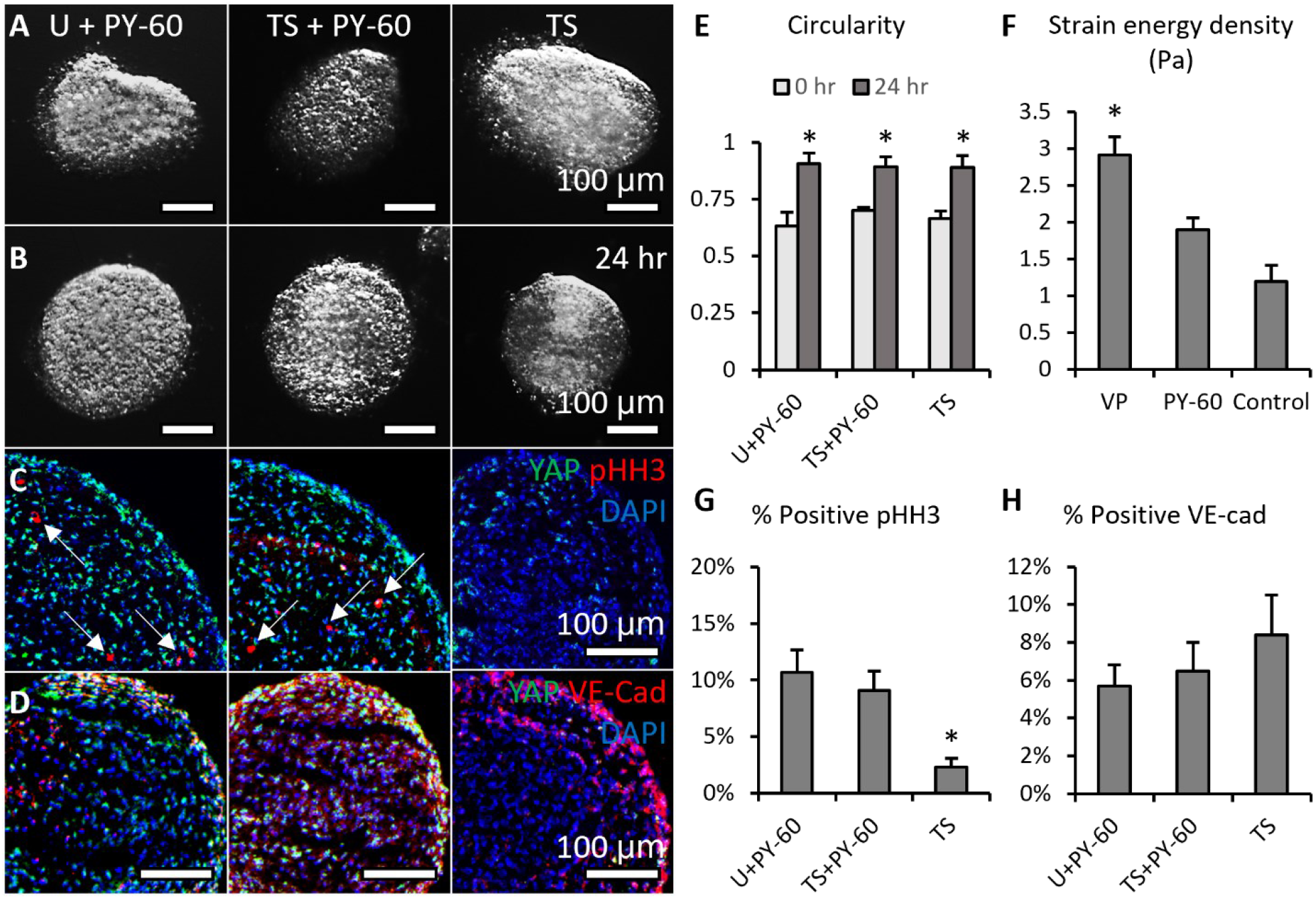
Activation of YAP promoted cell proliferation and tissue rounding. **A-B**. Bright field images of HH34 SL valve explants cultured under U+PY-60 (unloaded + YAP activator), TS+PY-60 (tensile stress + YAP activator), TS (tensile stress) conditions for 24 hours. **C**. After the 24-hr-culture, sections of explants were stained for YAP (green) and proliferation marker pHH3 (red, arrows), **D**. or YAP (green) and endothelial cell-cell junction VE-Cadherin (red). **E**. Comparison of explant morphology before and after 24 hour-culture. **F**. Micropipette aspiration measurement for the stiffness of valve explants cultured with YAP activator and inhibitor, compared with no treatment control. **G**. Comparison of ratios of cells expressing pHH3 under different culture conditions. **H**. Comparison of signal intensities of VE-Cad expression under different culture conditions, the intensities are normalized to maximum intensity. Data are presented by mean ± SEM, n = 15 explant valves from 8 embryos, *p < 0.05, two-way ANOVA tests or two-tailed student t-tests.

### YAP inhibition elevated valve stiffness

To assess the function of YAP in ECM stiffness, we employed micropipette aspiration to measure the strain energy density of the HH34 valve explants cultured with VP or PY-60 treatment. micropipette aspiration applies a local vacuum stress and monitors resultant tissue displacement within the tip (Figure S3). It has previously been used to measure material properties of chick embryonic valves during development.(34) Previous results showed the strain energy density between HH34 and HH36 valves increased more than four-fold. Here we found by inhibiting YAP, the strain energy density increased to almost three-fold of that of the freshly isolated valves (Figure 5H), close to the in-vivo stiffness increase. Although activation of YAP also led to an increase in strain energy density, it was not significant and far below the in-vivo stiffness increase. These results support that the stiffness of valves must increase with maturation in-vivo, which requires YAP inhibition.

### In-vivo mechanical manipulation led to valve defects and YAP mis-regulation

To manipulate mechanical forces in vivo, we performed LAL at early stages (HH24) when AV cushions were growing. We collected the hearts during cushion remodeling (HH31) when the valves begin to take shape. The LAL restricted the blood flow in the left ventricle, resulting in a reduced hemodynamic stress and shear stress on left AV valves but augmented forces on the right AV valves. As a result, the LAL valves and hearts (Figure 6A) had smaller sizes compared with the control valves and hearts (Figure 6B). Specifically, LAL led to an underdeveloped and globular left AV septal valve. While the right AV septal valve was overdeveloped and elongated. This is opposite to the normal valve development, during which the left AV valve has a much larger size and is more elongated (Figure 6C). LAL also caused an unbalanced YAP expression in left and right AV valves. We confirmed the same spatiotemporal pattern of YAP expression in chick embryonic valves as what we observed in mouse valves during equivalent stages (Figure S4). However, significantly fewer cells in the LAL left AV valve expressed YAP than the control (Figure 6C). Whereas a significantly higher ratio of YAP expressing cells was found in right AV valves.

**Figure 6.**
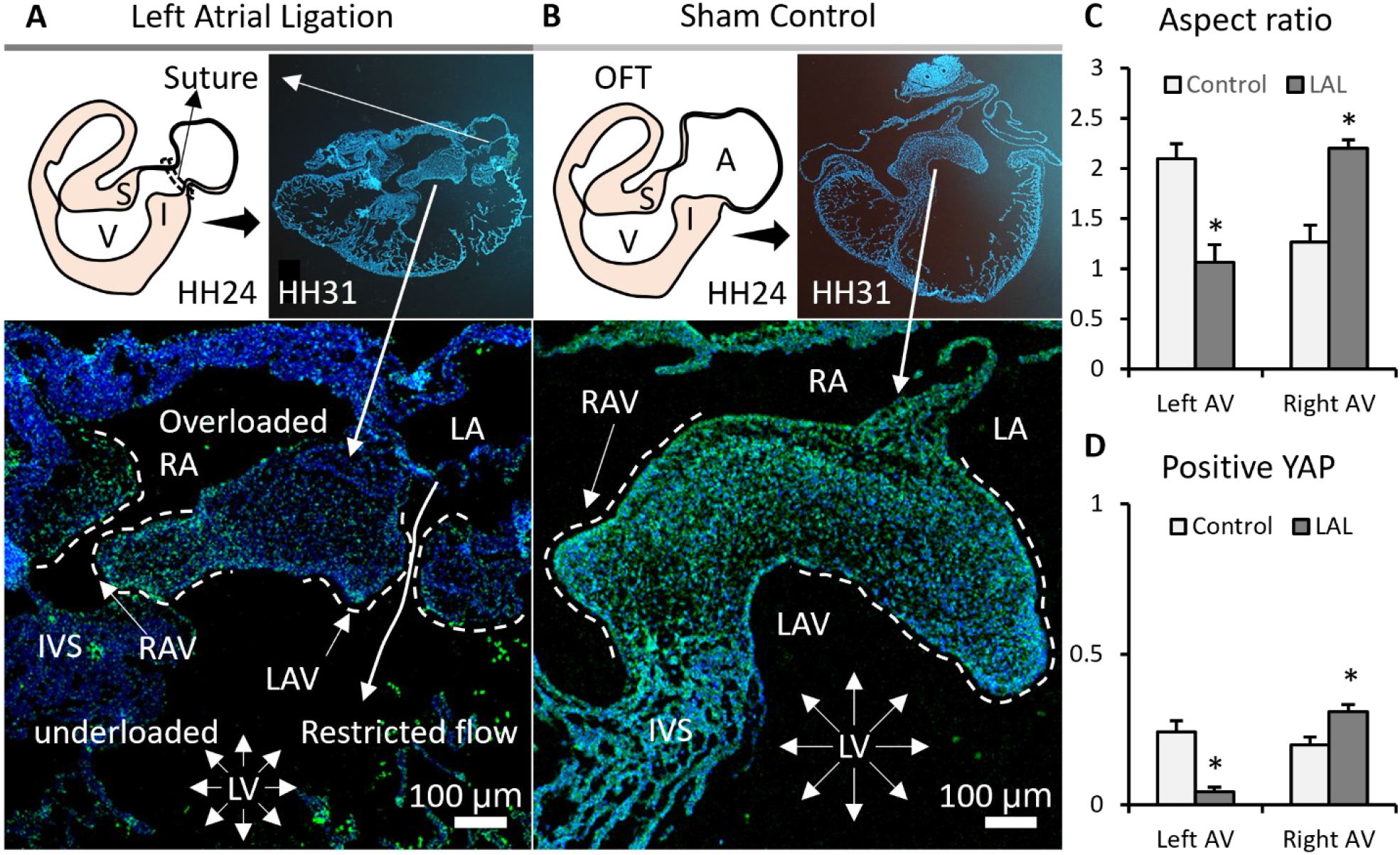
Interrupted biomechanics altered YAP activation and led to valve defects in-vivo. LAL performed at HH21 resulted in an underdeveloped left AV septal valve and overdeveloped right AV septal valve at HH31. **A**. Schematics of LAL experiments and YAP (green) straining of LAL heart sections from HH30 chick embryos. **B**. YAP (green) straining of sham control heart sections. **C**. Comparison of aspect ratios (which evaluate valve elongation) of AV septal valves between LAL and control. **D**. Ratios of cells expressing YAP in the AV septal valves. Data are presented by mean ± SEM, n = 6 sections from 3 embryos, *p < 0.05, two-tail student t-tests. A, atrium; V, ventricle; I, inferior cushion; S, superior cushion; LA, left atrium; RA, right atrium; LV, left ventricle; RV, right ventricle; IVS, interventricular septum; LAV, left atrioventricular septal valve; RAV, right atrioventricular septal valve.

## Discussion

Cardiac valves form in response to mechanical forces generated by the beating heart. These forces include shear and hydrostatic stress. Our results reveal that shear stress regulates valve shape by adjusting connections between VECs: USS (unidirectional shear stress) promotes an elongated shape by endothelium folding, while OSS (oscillatory shear stress) promotes a globular shape. Hydrostatic stress modulates valve size by controlling the proliferation of VICs: CS (compressive stress) promotes VIC proliferation and thus cushion growth while TS (tensile stress) inhibits cushion growth. It has been demonstrated that the interactions between mesenchymal and endothelial cells can form simple morphology.(35) Here we propose that a complex leaflet can also be achieved by a local collaboration between VECs and VICs in response to shear and hydrostatic stress. We use an ideal model (a symmetric cushion in a cylindrical tube) to elaborate this process (Figure 7A). The blood flow initiates an asymmetric deformation of the cushion (Figure 7B), leaving a gradient stress distribution with highest stress on the top.(36) A combination of OSS and CS drives a globular growth, while USS and CS work together to induce a flat and directional growth (Figure 7C). Due to the gradient stress distribution, CS localizes on the top even when the rest part of cushions is not compressed. The localized CS promotes a continuous local growth to fine tune the size and length (Figure 7D). USS tightens the inflow surface while OSS loosens the outflow surface. This generates a residual stress in the counterflow direction for the compaction of cushions. When the size and length are adequate, TS collaborates with USS to terminate growth (Figure 7E). Therefore, the spatiotemporally developed local forces ensure the proper size, length, and thickness of leaflets.

**Figure 7.**
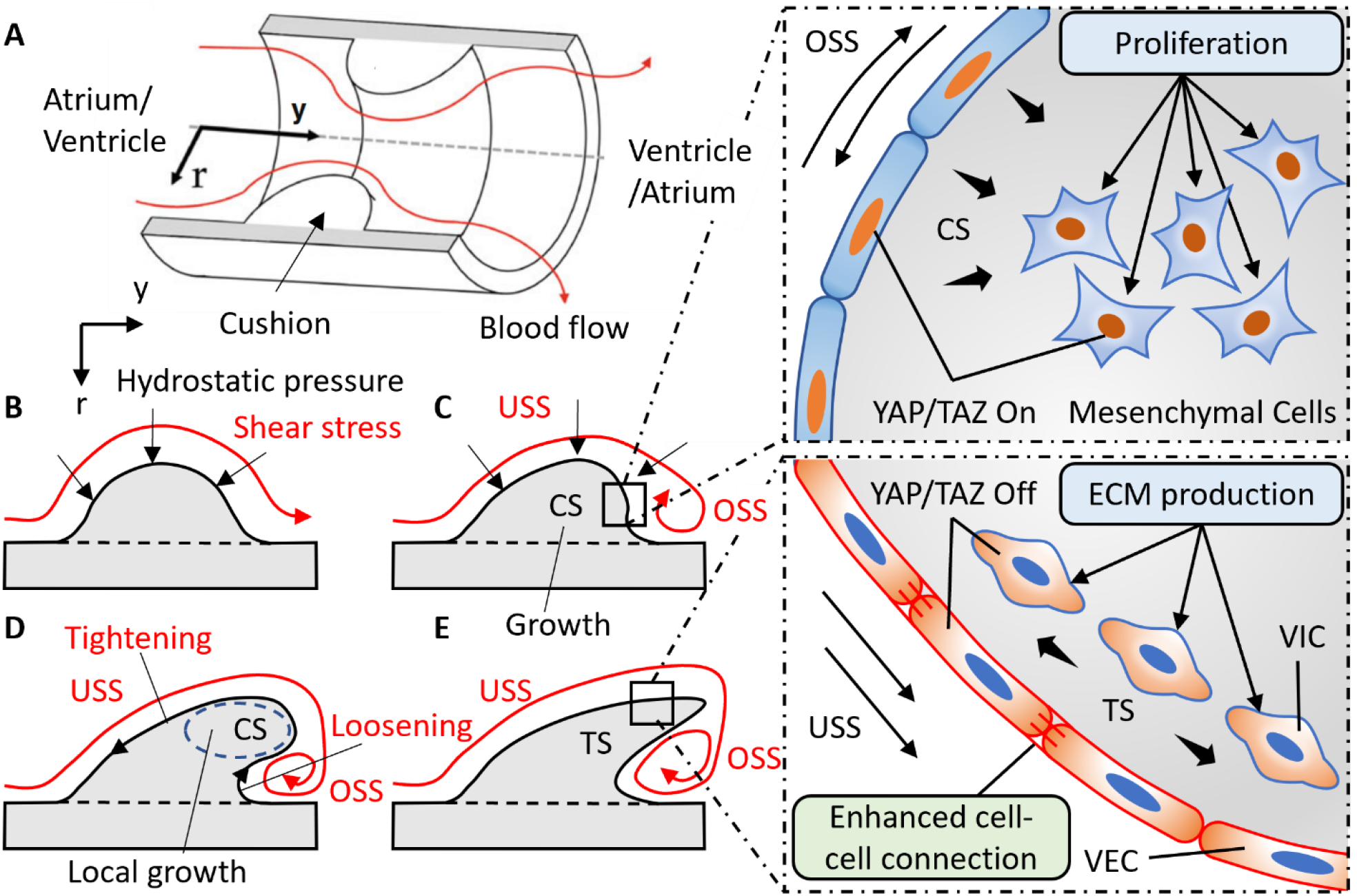
Local collaboration of shear and hydrostatic stress can guide complex morphogenesis via YAP signaling. **A**. A pseudo model of an endocardial cushion in the AV canal or OFT. **B**. Cross-section of the model shows the flowing blood generates hydrostatic stress and shear stress on cushions. **C**. CS promoted cushion growth by stimulating the proliferation of mesenchymal cells. OSS and CS activated YAP signaling in the VEC and VIC, respectively. **D**. Shear stress drives cushion remodeling into leaflet structure by modulating the cell-cell adhesions between VECs. **E**. The tensile stress (TS) and unidirectional shear stress (USS) inhibited YAP expression in the VEC and VIC, respectively.

On the other hand, disrupted flow with abnormally developed forces may cause errors in valve growth and remodeling. For example, a combination of delayed OSS and insufficient CS would result in a globular shape and small size, which are the features of underdeveloped valves. The delayed OSS and overloaded CS would cause a hypertrophic phenotype with globular shape and abnormally big size. Early occurrence of high USS and TS would promote a compacted and elongated shape but with an insufficient length.

This mechanism explains a major difference between contributions of cell lineage to the EMT and post-EMT growth and remodeling. During the EMT, cell lineage is a determinant factor. Cells derived from the second heart field make the major contribution to the OFT walls, and neural crest cells make a major contribution to the OFT cushions but a minor contribution to AV and intercalated cushions.(37, 38) However, during the post-EMT growth and remodeling, all valves with various morphologies are formed by identical VECs and VICs. Our study supports that the collaboration between VECs and VICs, instead of a specific cell lineage, determines the valve growth and remodeling.

To achieve the robust and precise control of valve growth and remodeling, local mechanosensing and response at the level of individual cells are required. YAP is a key mediator in mechanotransduction and a transcription cofactor of the Hippo pathway. During heart valve development, YAP is required for EMT,(39) but its role in post-EMT growth and remodeling is fully understood. Although knockout of VGLL4, another Hippo pathway cofactor that competes with YAP for DNA binding factors, led to malformations of aortic and pulmonary valves in mice,(7) the VGLL4 was only expressed in VICs and did not impact AV valves. This suggests that regulation of YAP is not solely dependent on VGLL4. Here we showed that YAP is robustly regulated by shear and hydrostatic stress and plays the same role in both OFT and AV valves growth and remodeling. The OSS promoted YAP activation in VEC, while USS inhibited YAP signaling. In the VIC, YAP nuclear translocation was boosted by CS but undermined by TS. Functionally, the YAP activation in VEC modulated cell-cell connections to strengthen or relieve surface tensions. The nuclear YAP raised the VIC proliferation rate.

These functions explain the spatiotemporal expression of YAP in the embryonic heart valves. At the early stage (E11.5), low stress OSS is the predominant force in the AV canal, which promotes YAP nuclear localization in the VEC (Figure 1A). It facilitates the dissociation of cell-cell connections and promotes the EMT process. As the EMT subsides, the cushions continue to grow. This leads to a constriction of the lumen, resulting in a higher shear stress that represses YAP activation in the VEC at E14.5 (Figure 1B). The YAP inhibition in VEC not only stops the EMT, but also strengthens local cell-cell adhesions to relieve surface tension so that the valve can elongate and compact. Meanwhile, due to the globular morphology, both valves are generally compressed at E14.5 under the hydrostatic stress generated by the flowing blood. This leads to a wide expression of YAP in the VIC, which promotes VIC proliferation and valve growth. During later stages (from E14.5 to E17.5), valves elongate into thin leaflets. The unidirectional blood flowing around the thin leaflets creates a YAP-suppressing USS on the inflow side and a YAP-inducing OSS on the outflow side (Figure 3A), resulting in the side specific pattern of YAP expression in VEC (Figure 1C). This delicate expression ensures a continued elongation until a mechanically sustainable size is reached. During this stage, as the elongated leaflets are stretched along flow direction, the tensile stress gradually overtakes compress stress in the elongated region. Only the bulged tip regions are still compressed by surrounding flow (Figure 3B), leading to a localization of YAP expression in the tip of valves (Figure 1C).

The mechanically induced heart valve defects are also associated with abnormal YAP expression. The LAL surgery diverted blood flow from the constricted left ventricle toward the untreated right ventricle, decreasing the hemodynamic stress on the left ventricle wall and cushions. This decreased stress undermines the CS generation in left AV cushions to suppress YAP expression, leading to an insufficient growth. Furthermore, according to the Law of Laplace, the decreased stress also results in a diminished ventricle wall tension that delays the transition from CS to TS, thus hindering cushion compaction. The LAL also altered shear stress environments. It has been demonstrated by computational fluid dynamics simulation that with LAL, the mean WSS applied on superior cushions decreased by 40%, and over 50% on inferior cushions.(26) The mean oscillatory shear index (which describes the level and duration of OSS) of left ventricle also increased significantly with LAL.(27) The delayed emerging of high stress USS could also contribute to the underdeveloped phenotype of the left AV by forcing the activation of YAP in the VEC to reinforce the spherical shape. By contrast, the OSS to USS transition was not significantly affected in the right ventricle, and the right AV cushion compacted and elongated as normal.

In conclusion, our study shows that the spatiotemporally coordinated mechanotransduction of shear and hydrostatic stress is required for proper valve growth and remodeling. The shear and hydrostatic stress can regulate VEC tensions and VIC proliferation by YAP pathways and thus determine the valve size and shape. Malfunctional YAP signaling could cause valve malformation, but improper local mechanical signaling imposes a more important malformation risk, even if the YAP signaling is fully functional and no genes are mutated. The presented mechanobiological system could also open an opportunity to control valve growth and change valve shape by influencing forces or targeting YAP pathway at a specific stage.

## Materials and Methods

### Shear stress bioreactor system

The bioreactor included a histology microscope slide, biocompatible double-sided tape (W.W. Grainger), 5 mm-thick silicone sheet (McMaster-Carr) and a sticky-Slide I Luer (Ibidi). Wells with 4mm diameters were created in the silicone sheet with a 4 mm disposable biopsy punch (Miltex). The magnitude of shear stress is determined by the height of the sticky-Slide I Luer and have been calculated and validated by Ibidi. The sticky-slide I Luer creates a 5 mm x 50 mm channel with various heights. 0.8 mm and 0.4 mm high sticky-slide I Luer were used to create shear stresses of 2 and 20 dyne/cm^2^, respectively. The components were clamped together with binder clips. The female Luers of the channel slide were connected to the silicone tubing with a 3.2 mm inner diameter (Size 16, Cole-Parmer).

For unidirectional shear stress (USS) experiments, flow was generated using Masterflex L/S Brushless variable-speed digital drive; Masterflex L/S 8-channel, 4-roller cartridge pump head; and Masterflex L/S large cartridges (Cole-Parmer). The peristaltic pump was connected to a pulse dampener (Cole-Parmer) to maintain non-pulsatile unidirectional flow over the samples. The pulse dampener was then connected to the bioreactors. The media was stored in a polycarbonate bottle with a filling/venting cap (Nalge Nunc International) with 80mL of M199 culture medium, with 1% insulin-selenium-transferrin, Pen/Strep, and 3% chick serum. The cells were exposed to a flow rate of 21.1 mL/min for 24 hours at 37°C and 5% CO2 within an incubator.

For oscillatory shear stress (OSS) experiments, flow was generated using a NE-1000 syringe pump (New Era Pump Systems, Inc.). The bioreactors were connected to a 20-mL syringe (BD Biosciences) that was controlled by the syringe pump. Cells were exposed to shear stress in the forward direction for one-half of a one-second cycle and in the reverse direction for the other half of the cycle at a flow rate of 21.1 mL/min for 24 hours at 37°C and 5% CO2 within an incubator.

### 3D endocardial cell culture

Collagen gels at a concentration of 2 mg/mL collagen were made using 3x Dulbecco’s Modified Eagle’s Medium (Life Technologies), 10% chick serum (Life Technologies), sterile 18 MΩ water, 0.1 M NaOH, and rat tail collagen I (BD Biosciences). An aliquot of the collagen gel solution was pipetted into the wells in the silicone sheet and allowed to solidify for 1 hour at 37°C and 5% CO2. The dissected outflow tracts were then placed on top of the collagen gel, and excess media was pipetted off to allow for the valve primordia to come in contact with the collagen gel. After 6 hours of incubation at 37°C and 5% CO2., the valve endocardial cells are repolarized, delaminated, and attached to the surface of the collagen constructs. These endocardial cells were then exposed to USS or OSS at 2 or 20 dyne/cm^2^ for 24 hours.

### Avian and AV cushion isolation and hanging drop culture system

Atrioventricular cushions (HH25) were dissected from the myocardium of embryonic chick hearts. The explants were cultured in M199 culture medium, 3% chick serum, and 1% insulin-transferrin-selenium. YAP inhibitor verteporfin (10 ug/ml) were purchased from Sigma Aldrich. The explants were placed in 20 ul hanging drops, settled at the apex of the droplets and cultured upside down for 24 hours.

### Left atrial ligation

Fertilized White Leghorn chicken eggs were incubated in a 38 °C forced-draft incubator to Hamburger-Hamilton (HH) stage 21 (3.5 days, Hamburger and Hamilton). The embryo was cultured in an ex-vivo platform previously described [RG 49]. Briefly, an overhand knot of 10-0 nylon suture loop was placed across a portion of either the right atrium and tightened, partially constricting the left AV orifice. This diverted flow from the constricted inlet toward the untreated inlet, decreasing hemodynamic load on the one side and increasing it on the other [RG 31, 32]. At D7 and D10 (HH31 and HH36, respectively), hearts and/or AVs were fixed, and paraffin sectioned for immunohistochemistry.

### Immunostaining

Chick and mouse hearts and mouse embryos were fixed in 4% paraformaldehyde for overnight, washed with TBS, embedded with paraffin, and sectioned. Sections were deparaffinized and rehydrated. Antigen retrieval was completed using citrate buffer at pH 6.0. Samples were then washed with TBS, permeabilized with 0.3% Triton-X 100 in TBS, and blocked with 3% BSA, 20mM MgCl, 0.3% Tween 20, 0.3M Glycine, and 5% Donkey serum in 1xTBS. Samples were then incubated with the primary antibodies at 1:100 dilution with the blocking solution overnight. Primary antibodies used include YAP (mouse DSHB YAP1 8J19, mouse Santa Cruz sc-101199, rabbit cell signaling #14074), Lef1 (rabbit Cell Signaling #2230), pHH3 (rabbit Cell Signaling #9701), VE-cad (abcam ab33168), MF-20 (mouse DSHB MF 20), IB4 (Vector B-1205-.5), Phalloidin (Invitrogen A12379). Samples were washed with 0.3% Triton-X 100 in TBS and incubated with secondary antibodies at 1:100 dilution with 5% BSA in TBS at room temperature for 1 hour. Secondary antibodies used include species-specific Alexa Fluor 568 or 647. Samples were then washed again for 3×10 minutes with TBS and stained with DAPI. Images were taken with Zeiss LSM 710 confocal microscope.

### Micropipette aspiration experiments

Mechanical properties of valve explants were measured by micropipette aspiration. A glass micropipette (rp&x2265;35 μm) was placed adjacent to the cushion surface collinear with the AV canal axis. Vacuum stress was incrementally applied via a 200 μL pipette calibrated with a custom manometer. Previous strain history was mitigated by preconditioning with 20 cycles of low pressurization (<1 Pa). The preconditioning step ensured the tissue and pipette tip were in full contact. Incremental stress loads were then applied, at which images were captured for each static load at 150X magnification using a Zeiss Discovery v20 stereo microscope. The aspirated length was measured using calibrated images in NIH ImageJ. An experimental “stretch ratio”, λ=(L+r_p_)/r_p_ was defined by normalizing the aspirated length to the pipette radius. The experiment stretch ratio is a measure of geometry change during aspiration, which is related, but not identical to the local stretch of the tissue. The ΔP versus λ curves were presented.

### Statistics

Images were analyzed using ImageJ. Results are presented as mean ± SD and compared using either an ANOVA tests with Tukey post hoc paired or two-tailed student t-tests. Differences were considered significant at p ≤ 0.05. For immunofluorescence staining, heart sections from n ≥5 embryonic hearts were used for quantification. For valve explants culture, n ≥5 independent cultures per treatment condition and four to six dozen chick embryos pooled for each experiment. For shear flow experiments, n ≥5 endocardial patches per shear flow condition. For LAL surgery, n ≥3 survival embryos for LAL control and sham control.

## Acknowledgements

This work was supported by NIH (grants HL128745, HL143247, HL160028), American Heart Association (grant 821615 to M.W.) and by biotechnology center via NYSTEM C029155 and NIH S10OD018516.

---

**Figure S1.**
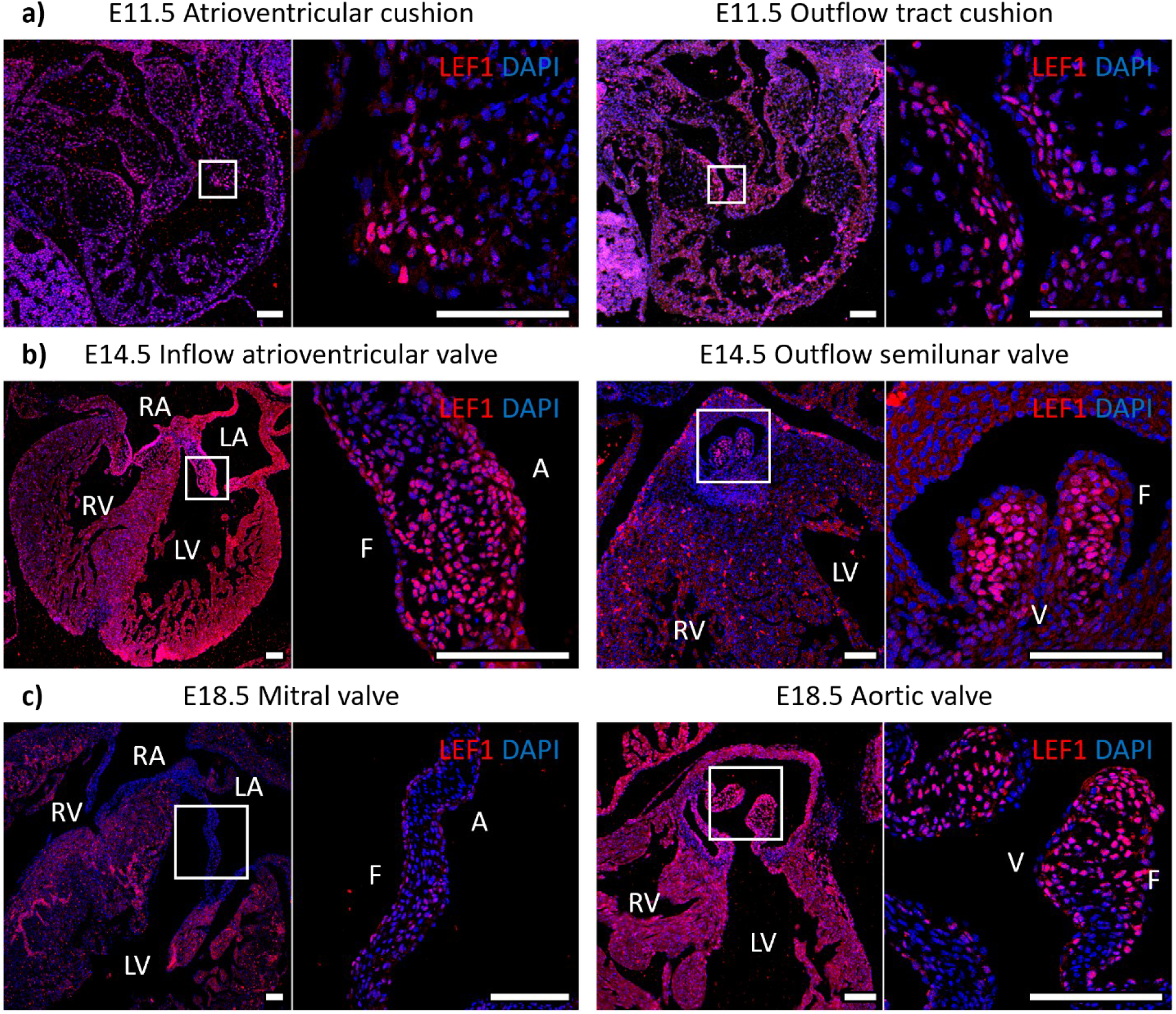
LEF1 expression at E11.5, E14.5 and E17.5 embryonic cardiac development. LA, left atrium; RA, right atrium; LV, left ventricle; RV, right ventricle; V, ventricularis; A, atrialis; F, fibrosa. Scale Bar = 100μm.

**Figure S2.**
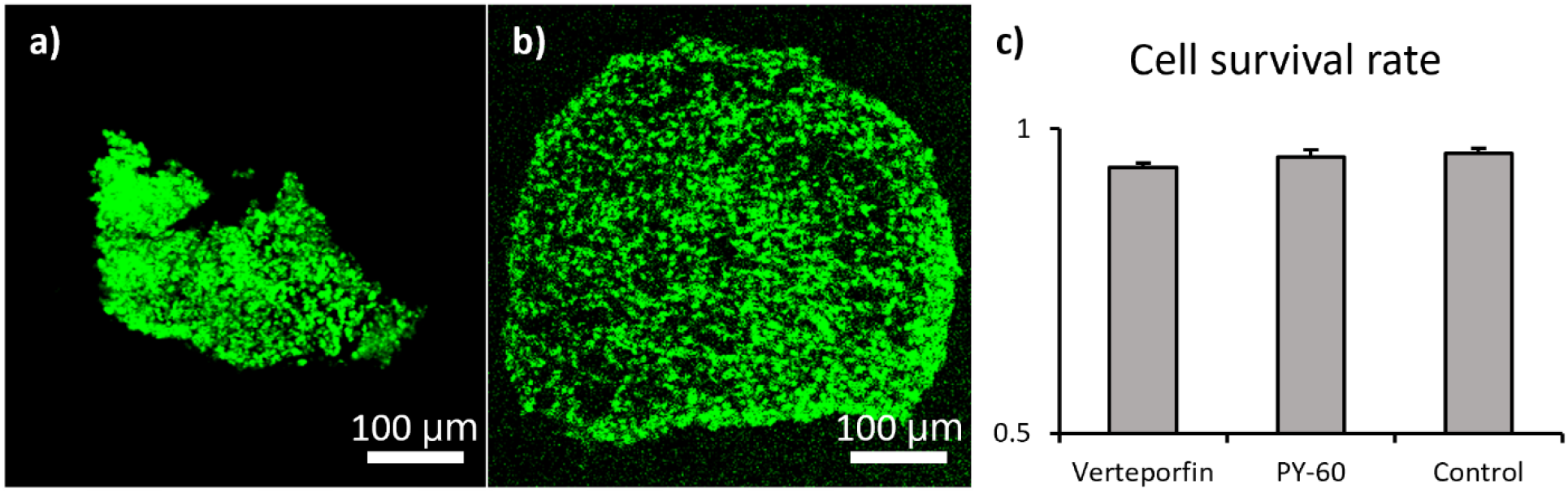
Verteporfin and PY-60 did not influence cell survival. Live and dead staining of **a)** verteporfin and **b)** PY-60 treated valve explants. **c)** Survival rates of cells in those valve explants compared with no pharmacological treatment control.

**Figure S3.**
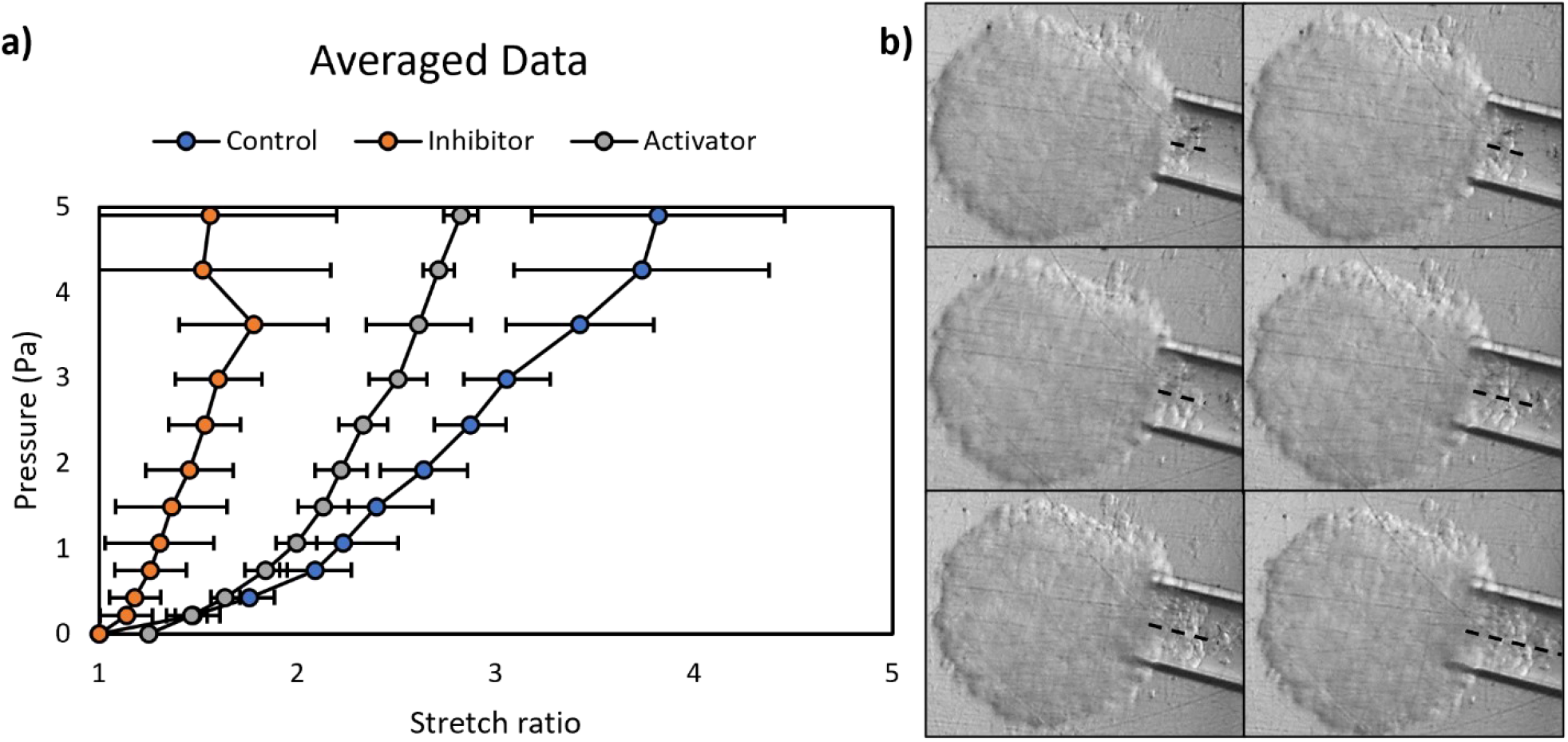
The stiffness of valve explants increased by YAP inhibition. **a)** Micropipette aspiration measurements of strain energy density for valve explants treated with YAP inhibitor and activator, and no treatment control. n ≥10 explants. **b)** Bright field images of the micropipette aspiration. Dash lines indicate tissue displacements within the tips.

**Figure S4.**
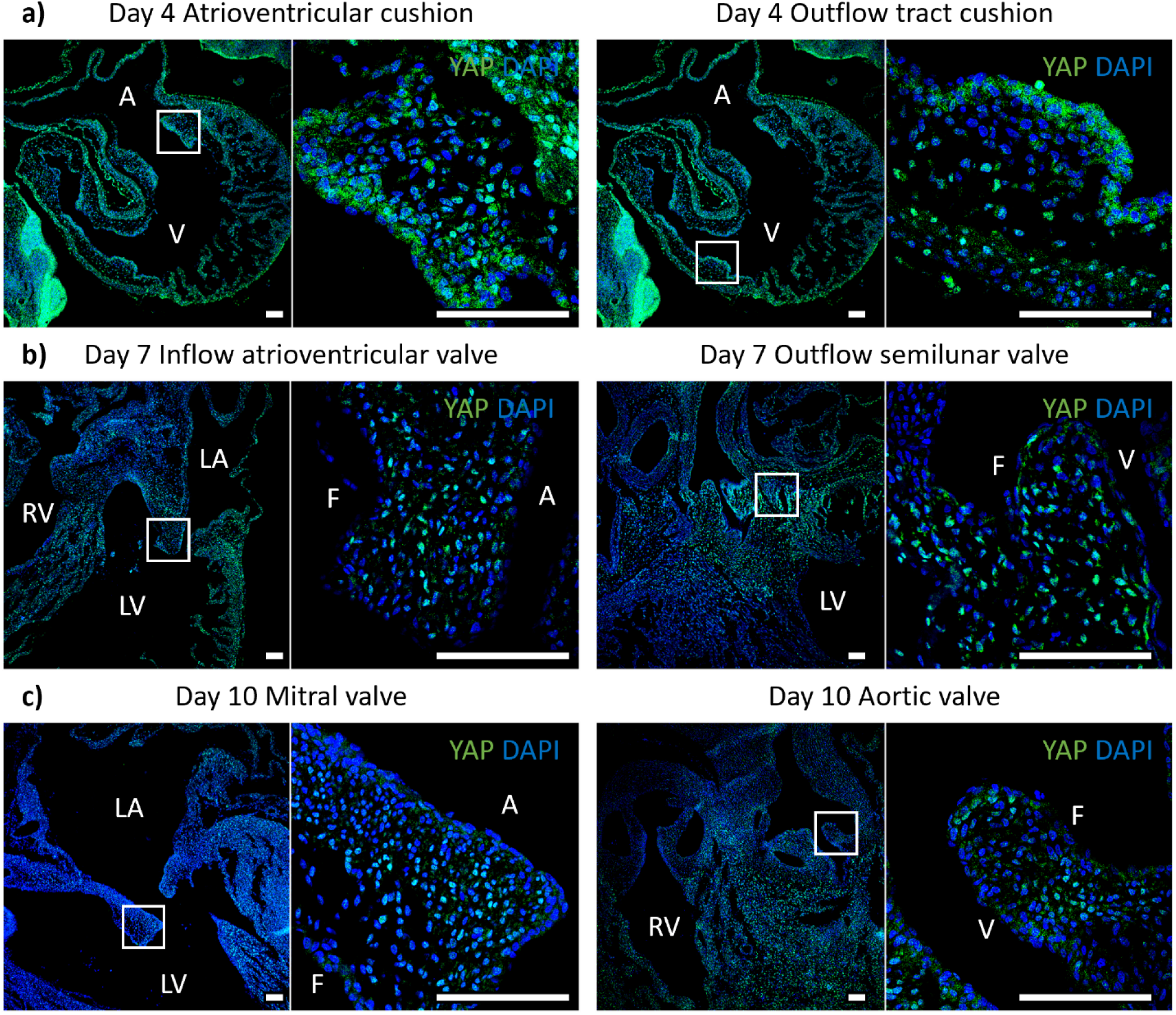
YAP expression in chick heart sections from Day 4, 7 and 10 during embryonic development. LA, left atrium; RA, right atrium; LV, left ventricle; RV, right ventricle; V, ventricularis; A, atrialis; F, fibrosa. Scale Bar = 100μm.

